# Genome-wide transcription during early wheat meiosis is independent of synapsis, ploidy level and the *Ph1* locus

**DOI:** 10.1101/437921

**Authors:** A.C. Martín, P. Borrill, J. Higgins, A.K. Alabdullah, R.H. RamÍrez-González, D. Swarbreck, C. Uauy, P. Shaw, G. Moore

## Abstract

Polyploidization is a fundamental process in plant evolution. One of the biggest challenges faced by a new polyploid is meiosis, particularly discriminating between multiple related chromosomes so that only homologous chromosomes synapse and recombine to ensure regular chromosome segregation and balanced gametes. Despite its large genome size, high DNA repetitive content and similarity between homoeologous chromosomes, hexaploid wheat completes meiosis in a shorter period than diploid species with a much smaller genome. Therefore, during wheat meiosis, mechanisms additional to the classical model based on DNA sequence homology, must facilitate more efficient homologous recognition. One such mechanism could involve exploitation of differences in chromosome structure between homologues and homoeologues at the onset of meiosis. In turn, these chromatin changes, can be expected to be linked to transcriptional gene activity. In this study, we present an extensive analysis of a large RNA-Seq data derived from six different genotypes: wheat, wheat-rye hybrids and newly synthesized octoploid triticale, both in the presence and absence of the *Ph1* locus. Plant material was collected at early prophase, at the transition leptotene-zygotene, when the telomere bouquet is forming and synapsis between homologues is beginning. The six genotypes exhibit different levels of synapsis and chromatin structure at this stage; therefore, recombination and consequently segregation, are also different. Unexpectedly, our study reveals that neither synapsis, whole genome duplication nor the absence of the *Ph1* locus are associated with major changes in gene expression levels during early meiotic prophase. Overall wheat transcription at this meiotic stage is therefore highly resilient to such alterations, even in the presence of major chromatin structural changes. This suggests that post-transcriptional and post-translational processes are likely to be more important. Thus, further studies will be required to reveal whether these observations are specific to wheat meiosis, and whether there are significant changes in post-transcriptional and post-translational modifications in wheat and other polyploid species associated with their polyploidisation.

## INTRODUCTION

Polyploidization, or whole genome duplication (WGD), has an important role in evolution and speciation, particularly in plants. It is now clear that all seed plants and angiosperms have experienced multiple rounds of WGD during their evolutionary history and are now considered to possess a paleopolyploid ancestry (Renny-Byfield et al., 2014). Polyploidy is traditionally classified into two separate types, autopolyploidy, arising from intraspecies genome duplication, and allopolyploidy, arising from interspecific hybridization. Many of the world’s most important crops, including wheat, rapeseed, sugarcane and cotton, are relatively recent allopolyploids; and much of the current knowledge about WGD is due to research involving these crop species. Several studies have reported major changes in transcription in somatic tissues following polyploidisation (Renny-Byfield et al., 2014 and references therein; Li et al., 2014; Edger et al., 2017). However, there have been very few previous reports on the effects of polyploidisation on transcription during meiosis, a critical stage in the establishment of a polyploid (Braynen et al., 2017).

Meiosis is the specialised cell division that generates haploid gametes for sexual reproduction. During meiosis, homologous (identical) chromosomes synapse along their length and recombine, leading to novel combinations of parental alleles, and ensuring proper chromosome segregation. Restriction of synapsis and crossover (CO) formation to homologous chromosomes (homologues) is therefore a prerequisite for regular meiosis. Subsequent recombination is also critical, not only to generate new combinations of genes, but also to ensure an equal distribution of genetic material and maintain fertility and genome stability across generations. One of the problems of polyploidisation is that it is initially accompanied by irregular meiosis, due to the presence of more than two identical homologues in autopolyploids, or very similar chromosomes (homoeologues) in allopolyploids. Thus, one of the biggest challenges faced by a new polyploid, is how to manage the correct recognition, synapsis, recombination and segregation of its multiple related chromosomes during meiosis, to produce balanced gametes.

Although studies on diploid model systems (reviewed in Mercier et al., 2016) have revealed much about the processes of recombination and synapsis, the way in which homologue recognition initiates the synapsis process during the telomere bouquet remains one of the most elusive questions still to be addressed. There are several different genetic and structural mechanisms of meiotic chromosome recognition reported in plants, mammals and fungi, indicating a differing process of recognition within different organisms (revised in Grusz et al., 2017). In most eukaryotes, homologous recognition is initiated by the formation of double-strand breaks (DSB) catalysed by the Spo11 protein. Subsequently, the DSB free ends invade the corresponding homologue regions, checking for sequence homology based on DNA sequence. However, it has also been observed, for example in hexaploid wheat, that the process of homologue recognition is also associated with major changes in chromosome chromatin structure (Prieto et al., 2004), suggesting that changes in chromatin structure may also be involved in the homologue recognition process. This may be more important in polyploid species such as hexaploid wheat, where the process of recognition must distinguish homologues from homoeologues. Hexaploid wheat *T. aestivum*, (2*n* = 6x = 42, AABBDD), also known as bread wheat, is a relatively recent allopolyploid, with three related ancestral genomes, which although different, possess a very similar gene order and content. Hexaploid wheat has a 16 Gb genome size, with high similarity between homoeologous genomes in the coding sequences (95-99%), and with a large proportion of repetitive DNA (11 85%) (IWGSC, 2018). Despite this, and the problem of having to distinguish between related chromosomes, hexaploid wheat is able to complete meiosis in a shorter period than diploid species such as rye, barley or even *Arabidopsis*, which possess a much smaller genome (Bennet et al., 1971; Bennet and Finch, 1971; Armstrong et al., 2003). Therefore, wheat meiosis is likely to exploit other mechanisms, apart from the traditional model based on DNA sequence homology, to facilitate homologous recognition. One such mechanism is likely to involve exploiting meiotic chromosome organisation, which in turn might be linked to the transcriptional activity of the genes on the homologous and homoeologous chromosomes (Cook, 1997; Wilson et al., 2005; Xu and Cook, 2008). It would be very interesting to assess the overall level of transcription occurring at the meiotic stage when chromosomes are recognising each other, and synapsis is beginning, to address whether the homology search influences or is influenced by transcription.

Despite the significant similarity between homoeologues, wheat behaves as a diploid during meiosis, with every chromosome recombining only with its true homologue. This phenotypic behaviour has been predominantly attributed to *Ph1* (*Pairing homoeologous 1*), a dominant locus on chromosome 5B (Riley and Chapman, 1958; Sears and Okomoto, 1958), which most likely arose during wheat polyploidisation (Chapman and Riley, 1970). In the absence of this locus, CO between non-homologues can occur, and so it was believed that the *Ph1* locus prevented synapsis between homoeologues. However, it has recently been demonstrated in wheat-wild relative hybrids lacking homologues, that although homoeologous chromosomes fail to synapse during the telomere bouquet (leptotene-zygotene transition), they do synapse to the same level after the telomere bouquet has dispersed, whether or not *Ph1* is present (Martín et al., 2017). This confirms that the *Ph1* locus itself does not prevent homoeologous synapsis after telomere bouquet dispersal in the wheat-wild relative hybrid. Similarly, in normal hexaploid wheat, only homologous synapsis can occur during the telomere bouquet stage. However, in the absence of *Ph1*, homologous synapsis is less efficient, with more overall synapsis occurring after the telomere bouquet has dispersed, when homoeologous synapsis can also take place. This non-specific synapsis between homoeologues leads to the low level of multivalents and univalents observed at metaphase I in wheat lacking *Ph1*. These observations indicate that, during the telomere bouquet, meiocytes from wheat and wheat-rye hybrids, with and without *Ph1*, exhibit major differences in level of synapsis, and chromatin structure. Such meiocytes provide a good source of material to assess the relationship between homologue recognition and synapsis, and transcription.

The *Ph1* locus was recently defined to a region on chromosome 5B containing a duplicated 3B chromosome segment carrying the major meiotic gene *ZIP4* and a heterochromatin tandem repeat block, inserted within a cluster of *CDK2-like* genes (Griffiths et al., 2006; Al-Kaff et.al., 2008; Martín et al., 2014, 2017). The duplicated *ZIP4* gene (*TaZIP4-B2*) within this cluster is responsible for both promotion of homologous CO and restriction of homoeologous CO, and is involved in improved synapsis efficiency (Rey et al., 2017, 2018a). The *CDK2-like* gene cluster has an effect on premeiotic events, its absence giving rise to delayed premeiotic replication and associated effects on chromatin and histone H1 phosphorylation (Greer et al., 2010). The processes of centromere pairing and telomere dynamics during premeiosis are also affected, probably as a result of this delay (Martínez-Pérez et al., 1999; Richards et al., 2012). Thus, the presence or absence of the *Ph1* locus affects the chromatin structure of chromosomes entering meiosis. This raises the question as to whether these premeiotic structural changes also affect overall transcription between homologues and homoeologues leading to altered recognition during early meiosis.

In this study we undertook a comprehensive study of transcription during early meiotic prophase, specifically at the leptotene-zygotene transition stage, when the telomere bouquet is formed in wheat and synapsis between homologues begins, to assess the effect on transcription of: changes in chromatin structure upon homologous recognition, level of synapsis, ploidy level, and presence of the *Ph1* locus. To evaluate this, a comparative transcriptome analysis was performed on meiocytes derived from wheat, wheat-rye hybrids and doubled wheat-rye hybrids (newly synthesized triticale), both in the presence and absence of *Ph1*. These six genotypes provided a unique set of transcription data, which can also be exploited in further studies. Surprisingly the analysis revealed that neither the level of synapsis, the ploidy level, nor the *Ph1* locus affected overall meiotic transcription during the leptotene-zygotene transition stage.

## MATERIALS AND METHODS

### Plant material

The plant material used in this study and its production is described in **Figure 1**, and includes: hexaploid wheat *Triticum aestivum cv*. Chinese Spring (2n = 6× = 42; AABBDD), either containing or lacking the *Ph1* locus (Sears, 1977); *rye Secale cereale* cv Petkus (2n = 2× = 14; RR); wheat-rye hybrids crosses between hexaploid wheat either containing or lacking the *Ph1* locus, and rye; octoploid triticale *× Triticosecale* Wittmack (2n = 8× = 56), obtained after genome duplication of wheat-rye hybrids either containing or lacking the *Ph1* locus.

**FIGURE 1.**
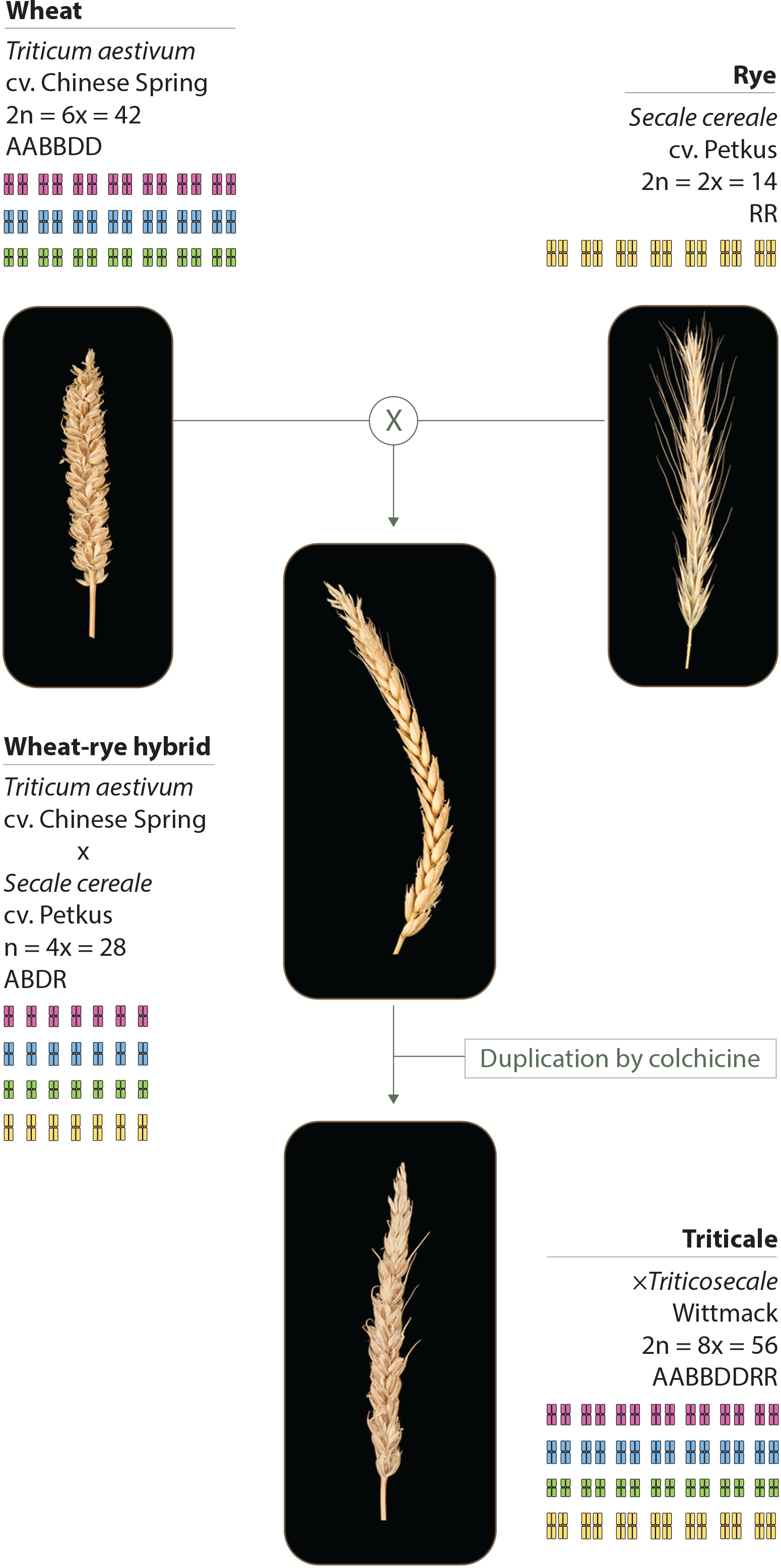
Plant material used in this study. Wheat (*T. aestivum* cv. Chinese Spring) was crossed as the female parent with rye (*S. cereale* cv. Petkus). The resulting interspecific wheat-rye hybrids are completely sterile. To obtain the octoploid triticale (× *Triticosecale* Wittmack), wheat-rye hybrids were treated with colchicine to double the chromosome number. The same crossing scheme was used to obtain the wheat-rye hybrids and the octoploid triticale, both lacking the *Ph1* locus; only this time, wheat lacking *Ph1* was used as the female parent for the initial crosses.

The wheat-rye hybrids were generated by either crossing *T. aestivum* cv. Chinese Spring or *T. aestivum* cv. Chinese Spring *Ph1*b mutant (Sears, 1977) as the female parent, with *S. cereale* cv. Petkus. Interspecific wheat-rye hybrids, either containing or lacking the *Ph1* locus, are completely sterile. The octoploid triticales were generated by treating wheat-rye hybrids, either containing or lacking the *Ph1* locus, with colchicine to double the chromosome number. Colchicine was applied according to the capping technique (Bell, 1950). Briefly, when the hybrids were at the 4-5 tillering stage, two of the tillers were cut and covered (or capped) with a small glass phial containing 0.5 ml of a solution of 0.3% colchicine. Once the solution was absorbed by the plant, the hybrids were left to grow. Successful chromosome doubling results in seed set. A few of the resulting duplicated seeds obtained were selfed twice in order to have sufficient seeds for all the studies. Only plants with a euploid chromosome number of 56 chromosomes were used for RNA-Seq sample collection. One spike of every plant used for the RNA-Seq analysis was selfed, so that cytological analysis could be performed on the progeny. One of the triticales lacking *Ph1* used for the RNA-Seq was sterile, so instead, another triticale lacking *Ph1* was used to complete the cytological analysis.

Seeds were germinated on Petri dishes for 3 to 4 days. The seedlings were vernalized for 3 weeks at 7 °C and then transferred to a controlled environment room until meiosis, under the following growth conditions: 16 h light/8 h night photoperiod at 20 °C day and 15°C night, with 70% humidity. After six to seven weeks, plants were ready for meiosis studies. Tillers were harvested after 6 to 7 weeks, at early booting stage, when the flag leaf ligule is just visible (39 Zadoks scale). Anthers were collected at early prophase, at the transition leptotene-zygote, which is during the telomere bouquet stage in wheat. For each dissected floret, one of the three synchronised anthers was squashed in 45% acetic acid in water to identify the meiotic specific stage. The two remaining anthers were harvested into RNAlater (Ambion, Austin, TX) for the RNA-Seq experiments or fixed in 100% ethanol/acetic acid 3:1 (v/v) for cytological analysis of meiocytes.

### Sample preparation and RNA extraction

Anthers from wheat, wheat-rye hybrids and triticale, both in the presence and absence of the *Ph1* locus, and rye were collected as described in the ‘Plant material’ section. Three biological replicates were prepared for each genotype, so a total of 21 samples were obtained. Anthers at the selected meiotic stage were harvested into RNA*later* (Ambion, Austin, TX). The anthers from three plants of each genotype were pooled in a 1.5-ml Eppendorf tube until 300 anthers were collected. As there were so few triticale seeds available, each triticale sample was derived from a single plant. Once sufficient anthers had been collected, the material was squashed using a pestle to release the meiocytes from the anthers, and the mix was transferred to a new Eppendorf trying to avoid as much of the anther debris as possible, to enrich the sample with meiocytes. The enriched meiocyte samples were centrifuged to eliminate the RNA*later* and homogenised using QIAshredder spin columns (Qiagen, Hilden, Germany). RNA extraction was performed using a miRNeasy Micro Kit (Qiagen, Hilden, Germany) according to the manufacturer’s instructions. This protocol allows purification of a separate miRNA-enriched fraction (used for further analysis) and the total RNA fraction (⍰200 nt) used in this study.

### RNA-Seq library preparation and sequencing

One microgram of RNA was purified to extract mRNA with a poly-A pull down using biotin beads. A total of 21 libraries were constructed using the NEXTflex™ Rapid Directional RNA-Seq Kit (Bioo Scientific Corporation, Austin, Texas, USA) with the NEXTflex™ DNA Barcodes– 48 (Bioo Scientific Corporation, Austin, Texas, USA) diluted to 6 μM. The library preparation involved an initial QC of the RNA using Qubit DNA (Life technologies, CA, Carlsbad) and RNA (Life technologies, CA, Carlsbad) assays, as well as a quality check using the PerkinElmer GX with the RNA assay (PerkinElmer Life and Analytical Sciences, Inc., Waltham, MA, USA). The constructed stranded RNA libraries were normalised and equimolar pooled into one final pool of 5.5 nM using elution buffer (Qiagen, Hilden, Germany, Hilden, Germany). The library pool was diluted to 2 nM with NaOH, and 5 μl was transferred into 995 μl HT1 (Illumina) to give a final concentration of 10 pM. Diluted library pool of 120 μl was then transferred into a 200-μl strip tube, spiked with 1% PhiX Control v3 and placed on ice before loading onto the Illumina cBot. The flow cell was clustered using HiSeq PE Cluster Kit v4, utilising the Illumina PE_HiSeq_Cluster_Kit_V4_cBot_recipe_V9.0 method on the Illumina cBot. Following the clustering procedure, the flow cell was loaded onto the Illumina HiSeq2500 instrument following the manufacturer’s instructions. The sequencing chemistry used was HiSeq SBS Kit v4 with HiSeq Control Software 2.2.58 and RTA 1.18.64. The library pool was run in a single lane for 125 cycles of each paired end read. Reads in bcl format were demultiplexed based on the 6 bp Illumina index by CASAVA 1.8, allowing for a one base-pair mismatch per library and converted to FASTQ format by bcl2fastq. RNA-Seq data processing: the raw reads were processed using SortMeRNA v2.0 (Kopylova et al., 2012) to remove rRNA reads. The non-rRNA reads were then trimmed using Trim Galore v0.4.1 (http://www.bioinformatics.babraham.ac.uk/projects/trim_galore/) to remove adaptor sequences and low-quality reads (-q 20—length 80—stringency 3).

### Chromosome coverage plots

RNA-Seq reads (trimmed nonrRNA reads as described above) for the 18 samples were aligned to the RefSeqv1.0 assembly (IWGSC, 2018), using HISAT v2.0.5 with strict mapping options (--no-discordant --no-mixed -k 1 --phred33 --rna-strandness RF) to reduce noise caused by reads mapping to the wrong regions. Output bam files (binary format for storing sequence alignment data) were sorted using samtools v1.5. The normalized average chromosome coverage (depth of reads aligning to each base along the genome) per 1 million base windows was obtained using bedtools v2.24.0. GenomeCoverageBed was run to generate a bedgraph file containing base coverage along each chromosome (scaled by reads per million). Each of the 21 chromosomes were divided into 1 million base windows and bedtools map was run to compute the average read depth over each 1 million base window. The average coverage was obtained for the three biological replicates of each sample. The ratio of coverage between samples was plotted as a heatmap using a custom R script (http://www.R-project.org). The following formula was used to calculate the ratio (−1 to +1) of coverage; ratio = sample1-sample2 / sample1+sample2.

### Differential expression analysis

Genes were examined for differential expression between wheat containing and wheat lacking the *Ph1* locus, by pseudoaligning the raw reads for these six samples against the Chinese Spring RefSeqv1.0+UTR transcriptome reference (IWGSC, 2018), using Kallisto v 0.42.3 (Bray et al. 2016) with default options. The index was built using a k-mer length of 31. Transcript abundance was obtained as estimated counts and transcripts per million (TPM) for each sample, and all samples were merged into matrices of gene-level expression using the script merge_kallisto_output_per_experiment_with_summary.rb from expVIP (https://github.com/homonecloco/expvip-web/blob/20180912ScriptToMergeKallistoOutput/bin/merge_kallisto_output_per_experiment_with_summary.rb). Only genes with a mean expression > 0.5 TPM in at least one condition, i.e. one of the genotypes (wheat containing or wheat lacking *Ph1*) were retained for differential expression analysis (Ramírez-González et al, 2018); this included 65,683 genes. Differential expression analysis between conditions (three replicates for each condition) was carried out using DESeq2 (v1.18.1) in R (v3.4.4). The numbers of differentially expressed genes at various thresholds (padj < 0.05, padj < 0.01 and padj < 0.001, and fold change > 2) were obtained. Gene Ontology (GO) enrichment was carried out using the R package goseq (v1.26.0) in R (v3.4.4). The GO term annotation was obtained from the RefSeqv1.0 (IWGSC, 2018). Genes up-regulated > 2-fold with padj < 0.01, and genes down-regulated > 2-fold with padj < 0.01 were analysed separately. Only significantly enriched GO terms (padj < 0.05) were retained. Human readable, PFAM and Interpro functional annotations for individual differentially expressed genes of interest were obtained from the functional annotation of the RefSeqv1.0 (IWGSC, 2018) from https://opendata.earlham.ac.uk/wheat/under_license/toronto/Ramirez-Gonzalez_etal_2018-06025-Transcriptome-Landscape/data/TablesForExploration/FunctionalAnnotation.rds(Ramírez-González et al., 2018).

Differential expression amongst wheat-rye hybrid and triticale samples was analysed by constructing an *in silico* wheat+rye transcriptome through combining the Chinese Spring RefSeqv1.0+UTR transcriptome reference (IWGSC, 2018) with the published rye transcriptome (Bauer et al., 2016). In total, this combined reference contained 326,851 transcripts, of which 299,067 were from wheat and 27,784 were from rye. The rye transcriptome only contained one isoform per gene whereas the wheat transcriptome contained multiple isoforms. The 12 samples with both wheat and rye genomes (wheat-rye hybrid and triticale in the presence and absence of *Ph1*, each with three biological replicates) were mapped to the *in silico* wheat-rye transcriptome using the same method as described above for wheat samples. Only genes with a mean expression > 0.5 TPM in at least one condition (wheat-rye hybrid or triticale, containing or lacking *Ph1*) were retained for differential expression analysis. This included 83,202 genes in total, of which 50,292 were high confidence (HC) wheat genes, 22,138 were low confidence (LC) wheat genes and 10,772 were rye genes. Two comparisons were carried out to examine the effect of *Ph1* (wheat-rye hybrids carrying vs lacking *Ph1*, and triticale carrying vs lacking *Ph1*), and one comparison performed to examine the effect of different synapsis levels and chromosome doubling (wheat-rye hybrids vs triticale, both carrying *Ph1*). Differentially expressed genes for each comparison were identified using DESeq2 as described above, using three replicates per condition. The numbers of differentially expressed genes at various thresholds (padj < 0.05, padj < 0.01 and padj < 0.001, and fold change > 2) were obtained for wheat and rye genes. GO term annotation was available for the wheat genes, therefore we focused only on differentially expressed wheat genes for GO enrichment analysis and excluded differentially expressed rye genes from this analysis. GO enrichment analysis was carried out as described for the wheat samples, again separately analysing genes up-regulated > 2-fold with padj < 0.01, and genes down-regulated > 2-fold with padj < 0.01.

The rye genome sequence is still incomplete; therefore, the present study focused on wheat-specific transcription effects. However, the rye data (PRJEB25586) are deposited to facilitate future analyses by the community.

### Genomic in situ hybridisation of mitotic and meiotic cells (GISH)

It was not possible to analyse the samples used for the RNA-Seq analysis, so their progeny were analysed instead. One spike of every plant used for the transcription analysis was selfed, and three individuals from each progeny analysed. One of the triticale lacking *Ph1* used for the RNA-Seq was sterile, so instead, another triticale lacking *Ph1* which had 57 chromosomes, was used.

The preparation of mitotic metaphase spreads and subsequent genomic in situ hybridisation (GISH) was carried out as described previously (Rey et al., 2018b). Meiotic metaphase I spread preparation and subsequent GISH were also carried out as described previously (Cabrera et al., 2002). *S. cereale, Triticum urartu* and *Aegilops taushii* were used as probes to label rye, wheat A- and wheat D-genomes respectively. *S. cereale* genomic DNA was labelled with tetramethyl-rhodamine-5-dUTP (Sigma) by nick translation as described previously (Cabrera et al., 2002). *T. urartu* and *Ae. taushii* genomic DNA were labelled with biotin-16-dUTP and digoxigenin-11-dUTP, using the Biotin-nick translation mix and the DIG-nick translation mix respectively (Sigma, St. Louis, MO, USA) according to the manufacturer’s instructions. Biotin-labelled probes were detected with Streptavidin-Cy5 (Thermo Fisher Scientific, Waltham, Massachusetts, USA). Digoxigenin-labelled probes were detected with anti-digoxigenin-fluorescein Fab fragments (Sigma).

Images were acquired using a Leica DM5500B microscope equipped with a Hamamatsu ORCA-FLASH4.0 camera and controlled by Leica LAS X software v2.0. Images were processed using Fiji (an implementation of ImageJ, a public domain program by W. Rasband available from http://rsb.info.nih.gov/ij/) and Adobe Photoshop CS4 (Adobe Systems Incorporated, USA) version 11.0 × 64.

### Availability of supporting data

Raw Illumina reads have been deposited into EMBL-EBI ENA (European Nucleotide Archive, https://www.ebi.ac.uk/ena" under project number PRJEB25586. Analysed data for the wheat samples (TPM and counts) were integrated in the expVIP platform www.wheat-expression.com (Borrill et al., 2016).

## RESULTS

### 1. Transcriptome sequencing

To assess whether synapsis, ploidy level and changes in chromatin structure associated with *Ph1* have any effect on global transcription during early meiotic prophase I, the transcriptome of wheat, wheat-rye hybrid and the corresponding triticale were analyzed by RNA-Seq in the presence and absence of the *Ph1* locus. In wheat florets, the three anthers and the meiocytes within them are highly synchronized in development. We staged one of the three anthers by microscopy, to ensure that the meiocytes were at the transition leptotene-zygotene stage, leaving the other two anthers for RNA extraction. Three biological replicates were produced for each transcriptome, with a total of 18 libraries generated.

Using Illumina sequencing, a total of 1,388 million reads were generated for the 18 libraries. For subsequent analysis, the RNA-Seq data were processed using two different methods. Firstly, samples were trimmed and aligned to the wheat RefSeqv1.0 assembly using HISAT to generate the chromosome coverage plots. Strict mapping options were used to reduce the noise caused by reads mapping to the wrong regions, particularly mis-mapping from the rye onto the wheat genome. The percentage of reads aligned to the wheat genome was on average 88.46% for wheat samples, 74.61% for wheat-rye hybrids samples and 75.48% for triticale samples (**Table S1**). On average 13.4 % fewer reads mapped in wheat-rye and triticale samples than in wheat samples, indicating that the stringent mapping conditions were effective in reducing mis-mapping. However, it is possible that a low level of residual mis-mapping occurred, whereby reads from rye genes are mapped onto the wheat genome.

Secondly, DESesq2 was used to examine genes differentially expressed between genotypes. The six wheat samples were pseudoaligned to the Chinese Spring RefSeqv1.0+UTR transcriptome reference using kallisto. The percentage of reads pseudoaligned was similar across samples, with a mean value of 72% as detailed in **Table S2**. The three biological replicates showed good correlation and clustered together in the principle component analysis, although samples containing the *Ph1* locus grouped together more tightly than samples lacking *Ph1* (**Figure S1A**). The wheat RefSeq Annotation v1.0 includes 110,790 HC genes and 158,793 LC genes. LC genes represent partially supported gene models, gene fragments and orphan genes (IWGSC, 2018). We decided to retain these LC genes in our differential expression analysis because one of the reasons for them to be considered as LC genes is a lack of RNA-Seq data evidence. As already mentioned, no data on wheat meiosis were previously published, and since it is therefore possible that some of these LC genes are specifically involved in meiosis, we decided to include them in the analysis.

The 12 samples from wheat-rye hybrids and triticale, were mapped to a wheat+rye transcriptome created *in silico* for this study (described in material and methods). Although our aim in these samples was principally to study the expression of wheat genes, the use of a hybrid transcriptome reduced the possibility of mis-mapping between rye and wheat reads, which would have led to inaccurate quantification of expression. Kallisto was used for mapping because in wheat, it accurately distinguishes reads from homoeologues carrying genes with a sequence identity between 95-97 % (Ramírez-González et al., 2018). Therefore, kallisto is capable of distinguishing wheat and rye reads, which are more divergent (91 % sequence identity within genes (Khalil et al., 2015)). The percentage of reads pseudoaligned to this transcriptome was similar across samples, with a mean value of 71% for wheat-rye hybrids and triticale (**Table S2**). The three biological replicates from each genotype showed good correlation and clustered together in the principle component analysis (**Figure S1B**). In total, 269,583 genes from wheat (110,790 HC and 158,793 LC), plus 27,784 genes from rye were annotated, giving a total of 297,367 genes for the hybrid transcriptome created.

### 2. Overall transcription is independent of synapsis, ploidy level or the presence of *Ph1*

Chromosome coverage plots were generated to reveal a global picture of the difference in transcription between the different genotypes analyzed. The cleaned RNA-Seq reads were aligned to the RefSeqv1.0 assembly and the ratio of the coverage along all the chromosomes plotted as a heatmap (**Figure 2**).

**FIGURE 2.**
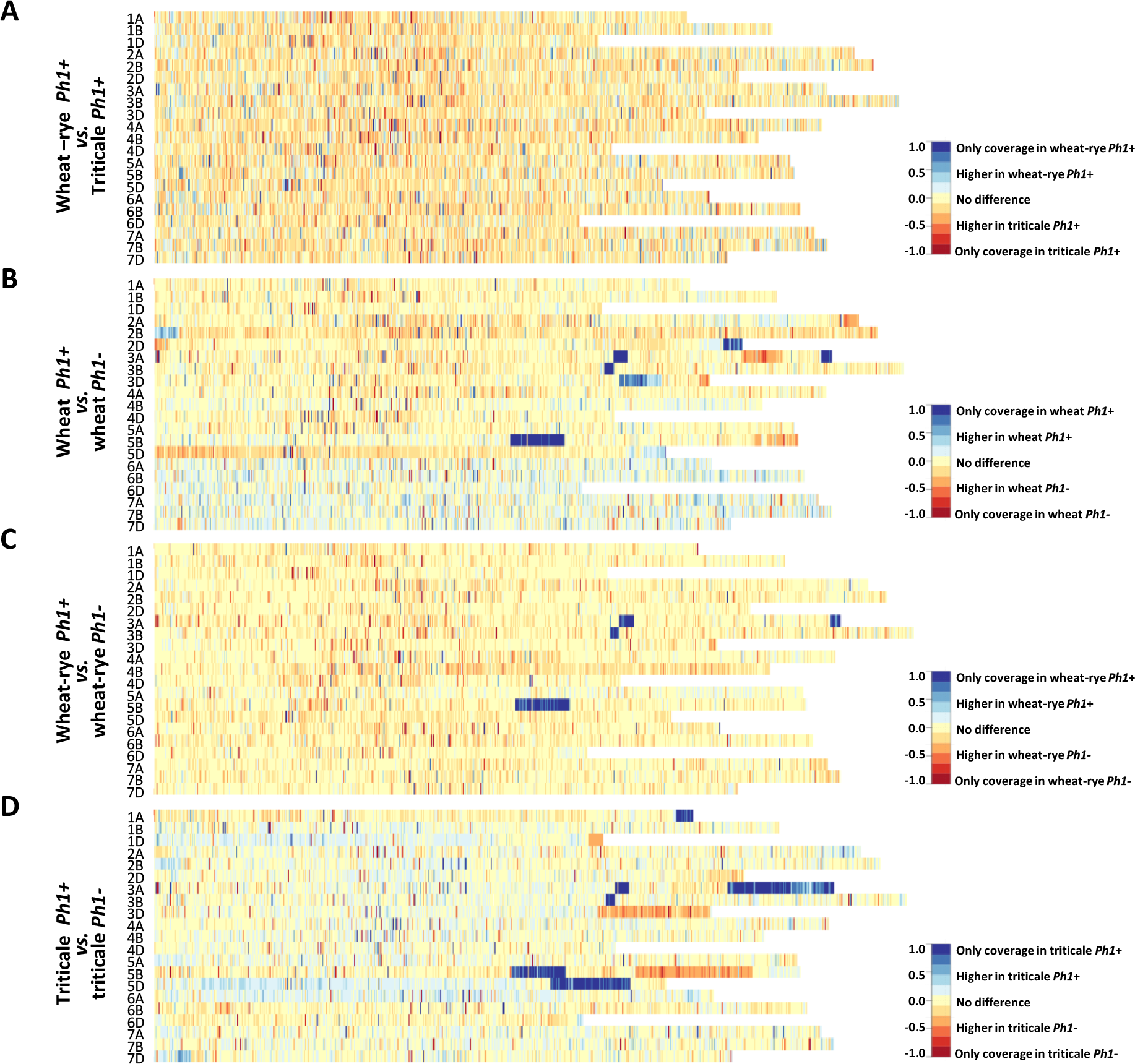
Chromosome coverage plots generated using the obtained RNA-Seq data. The ratio of coverage along all chromosomes was plotted as a heatmap. **(A)** Heatmap comparing wheat-rye hybrids and triticale, both containing the *Ph1* locus (*Ph1+*). No global change in transcription was observed between these two genotypes. **(B)** Heatmap comparing wheat in the presence (*Ph1+*) and absence (*Ph1*-) of *Ph1*. **(C)** Heatmap comparing wheat-rye in the presence and absence of *Ph1*. **(D)** Heatmap comparing triticale in the presence and absence of *Ph1*. Several deletions (visualized in dark blue) and other chromosomes reorganizations were detected in all genotypes in the absence of *Ph1* **(B-D)**.

At the leptotene-zygotene transition, when the telomere bouquet is tightly formed, only homologous chromosomes can synapse. In wheat-rye hybrids there are no homologues present, and therefore, no synapsis takes place at this stage; whereas in triticale, a significant level of synapsis occurs. However, no overall change in wheat transcription was observed when these two genotypes were compared (**Figure 2A**, **Figure S2**). Even more striking was that the duplication of the genome and change in ploidy level had little effect on overall wheat transcription. The wheat-rye hybrid is a poly-haploid (n = 4× = 28, ABDR) and triticale an octoploid (2n = 8× = 56, AABBDDRR), and although vegetative development is normal in both genotypes, the haploid hybrids are completely sterile. The reason for sterility in the wheat-rye hybrid is that meiosis is highly compromised, with only one CO at metaphase I and subsequent random segregation of chromosomes. Despite this, wheat transcription during early meiotic prophase seems not to be affected.

Next, a comparison of wheat, wheat-rye hybrid and triticale, with their corresponding genotypes lacking the *Ph1* locus was made (**Figure 2B-D**, **Figure S2**). This time, the heatmaps revealed a very different situation, with very clear differences in transcription in all comparisons. A deletion on chromosome 5B (visualized in dark blue in **Figure 2B-D**) was observed in all samples lacking *Ph1*, corresponding to the deletion of the *Ph1b* mutant (Sears, 1977). However, several other deletions (visualized in dark blue) were also observed in all samples. Due to the nature of this locus, which affects synapsis and CO formation between homoeologues, the presence of chromosome rearrangements has been previously described in wheat lacking *Ph1* (Sánchez-Morán et al., 2001). However, the number of rearrangements revealed was higher than expected. An interstitial deletion on 3BL and two deletions on 3AL, one interstitial and another terminal, were common to all samples lacking *Ph1*. Apart from these, wheat lacking *Ph1* had two more deletions: a terminal one on 2DL and a distal one on 3DL; while triticale lacking *Ph1* had three more deletions: a terminal one on 1AL, a large terminal one on 3AL and a large distal one on 5DL. As observed in **Figure 2C**, the heatmap belonging to the wheat-rye hybrids showed no difference in overall transcription whether *Ph1* was present or absent, apart from the common deletions mentioned. However, heatmaps corresponding to wheat and triticale were more difficult to interpret, with several chromosome regions showing clear differential expression without a completely clear-cut presence or absence of *Ph1*, as in the case of the deletions. The triticale heatmap (**Figure 2D**) for example, revealed three chromosome regions showing increased transcription in triticale lacking *Ph1* (visualized in orange in the heatmap): a terminal region in 1DL and 3DL, and a large distal region on 5BL. Interestingly, every one of these chromosome regions corresponded to a chromosome deletion on a homoeologous chromosome. For example, the terminal deletion on 1AL, corresponded to the terminal increased transcription on 1DL. Recombination could occur between 1A and 1D in the absence of *Ph1*, resulting in two 1D chromosomes plus two 1A chromosomes being detected in this material, in which the distal part of 1AL had a translocation from 1DL. This, therefore, resulted in four copies of the D-genome chromosome segment, and hence increased transcription. These observations are consistent with wheat nullitetrasomic line analysis (Borrill et al, 2016), where the presence of four copies of a homoeologue leads to a doubling of transcription. The observed increases in transcription associated with the other deletions in wheat and triticale lacking *Ph1* could also be explained in a similar manner. There were some regions where the interpretation was more complex. To investigate this further, we created heatmaps of wheat vs. wheat lacking *Ph1*, and triticale vs. triticale lacking *Ph1*, for every individual *Ph1* mutant sample (**Figure S3, Figure S4**). Results revealed that every individual sample lacking *Ph1* was different, apart from the rearrangements common to all samples lacking *Ph1* described above. One triticale sample was extremely rearranged (**Figure S4**), so we decided to explore this further and perform GISH experiments on this material, which will be described in the following sections.

In summary, at this stage of meiosis, overall transcription was not affected by the absence of the *Ph1* locus. Therefore, chromatin changes associated with the *Ph1* locus did not affect overall transcription. All significant transcriptional changes observed between genotypes with and without *Ph1* were associated with the presence/absence of chromosome regions likely to be the result of homoeologous recombination. We therefore conclude that neither synapsis, level of ploidy nor the presence of *Ph1* have a significant overall effect on wheat meiotic transcription.

### 3. Analysis of differentially expressed genes (DEG)

Chromosome coverage plots showed no global changes in transcription. However, we also wanted to identify the number of genes differentially expressed, and check whether they were related to meiotic processes. A Kallisto-DESeq2 pipeline was used to examine the DEG between genotypes. Only genes expressed > 0.5 TPM in at least one of the genotypes were selected for differential expression analysis, the rest being filtered out as non-expressed genes. The number of DEG among samples was calculated using different thresholds as described in material and methods and **Table S3**, with further analysis focussed on the comparisons at padj < 0.01 and fold change > 2.

#### 3.1 DEGs between wheat-rye hybrids and octoploid triticale (both containing the Ph1 locus)

Of 297,367 genes present in the wheat+rye hybrid transcriptome, 83,202 genes (27.98%) were expressed in our samples, of which 72,430 were from wheat and 10,772 from rye (**Table S4**). As the rye genome sequence is incomplete, and most genes have no functional annotation, the analysis only focused on wheat genes. The 72,430 wheat genes include both high-confidence (HC) and low-confidence (LC) genes (**Table 1**). Although both HC and LC genes were included in the analysis, results are presented for HC genes separately in **Table S4**. Interestingly, a high percentage of LC genes (22,138 genes) were detected as being expressed during early meiosis, with 2,711 LC genes expressed at a relatively high level (>10 TPM). This provides evidence that these LC genes could be HC genes which were missed during the original annotation process, perhaps due to the lack of RNA-Seq data from meiosis samples. Among the 72,430 wheat expressed genes, only 344 genes were differentially expressed between wheat-rye hybrids and triticale (Table 1). This means that DEGs represent only 0.47% of all genes, a strikingly low number considering that the whole genome has been duplicated. These results also indicate, consistent with the chromosome coverage plots, that overall gene expression is independent of synapsis and the absence of homologous chromosomes.

**TABLE I.** Annotated genes, which were differentially expressed between samples in this study. *Ph1+* = containing the *Ph1* locus, *Ph1* - = lacking the *Ph1* locus.

| Comparison | Up-regulated genes | Down-regulated genes | Total DEG | % DEG | Total expressed genes |
| --- | --- | --- | --- | --- | --- |
| Wheat-rye hybrid *Ph1+* vs. triticales *Ph1+* | 150 | 194 | 344 | **0.47** | 72,430 |
| Wheat-rye hybrid *Ph1+* vs. wheat-rye hybrid *Ph1-* | 478 | 95 | 573 | **0.79** |  |
| Triticales *Ph1+* vs. triticales *Ph1-* | 1,779 | 893 | 2,672 | **3.69** |  |
| Wheat *Ph1+* vs. wheat *Ph1-* | 450 | 24 | 474 | **0.72** | 65,583 |
| All *Ph1+* vs. All *Ph1-* | 353 | 5 | 358 | **0.56** | 63,529 |

To check whether DEGs genes were associated with common processes, Gene Ontology (GO) term enrichment was carried out on genes differentially expressed between samples (**Table S5**). Genes down-regulated upon chromosome doubling (up-regulated in the hybrids *vs.* triticale) were enriched for few GO terms, which were mostly related to metabolic processes (general functions) not related to meiosis; moreover, p-values were only just significant (0.02 to 0.05). In the case of genes up-regulated upon chromosome doubling (down-regulated in the hybrids *vs.* triticale), two thirds of the GO terms were related to stress and response to external stimuli, the rest being related to cell communication and catabolic processes. Although the present study focuses on changes in overall gene expression, we also extracted the functional annotation for all DEG, available at **Table S6**.

#### 3.2 DEGs in the absence of the Ph1 locus

Next, we identified DEGs between wheat, wheat-rye hybrid and triticale, and their corresponding samples in the absence of the *Ph1* locus. Of 269,583 genes annotated (RefSeqv1.0 assembly), 65,583 were expressed in our wheat samples, from which 474 genes (0.72%) were differentially expressed when *Ph1* was deleted (**Table 1**). In the case of wheat-rye hybrids, 573 wheat genes (0.79%) were differentially expressed in the absence of *Ph1*; and 2672 (3.69%) genes were differentially expressed for triticale lacking *Ph1* (**Table 1**). The number of DEGs was higher in triticale, in agreement with the chromosome coverage plot results. As described previously, all genotypes lacking *Ph1* exhibit deletions and other chromosome rearrangements, and the DEGs detected could therefore be a consequence of these reorganizations, rather than due to an absence of *Ph1* alone. To exclude these DEGs being a consequence of chromosomal rearrangement, we identified DEGs shared by all comparisons. We found that all three comparisons had 358 DEGs in common (**Figure 3**), of which 186 genes were located in the *Ph1b* deletion on 5B. A further 106 and 33 genes were located in deletions common to all genotypes lacking *Ph1* on 3A and 3B respectively. Therefore, in total there were only 33 DEGs which could not be accounted for, based on their location within a common deleted region, meaning that overall gene expression was not significantly altered during early prophase by the absence of the *Ph1* locus. We did not observe any trend directly related to meiosis in the functional annotation of these 33 genes (**Table S7**). One gene, TraesCS2A01G561600, annotated as a DNA/RNA helicase protein, could be potentially involved in meiosis, since these enzymes play essential roles in DNA replication, DNA repair and DNA recombination, which occur both in somatic and meiotic cells. However, the syntenic ortholog in *Arabidopsis*, the chromatin remodelling 24 gene (*CHR24/ AT5G63950*) is a member of the *SWI2/SNF2* family known to be involved in DNA repair and recombination in somatic tissue, while no function during meiosis has been reported (Shaked et al, 2006).

**FIGURE 3.**
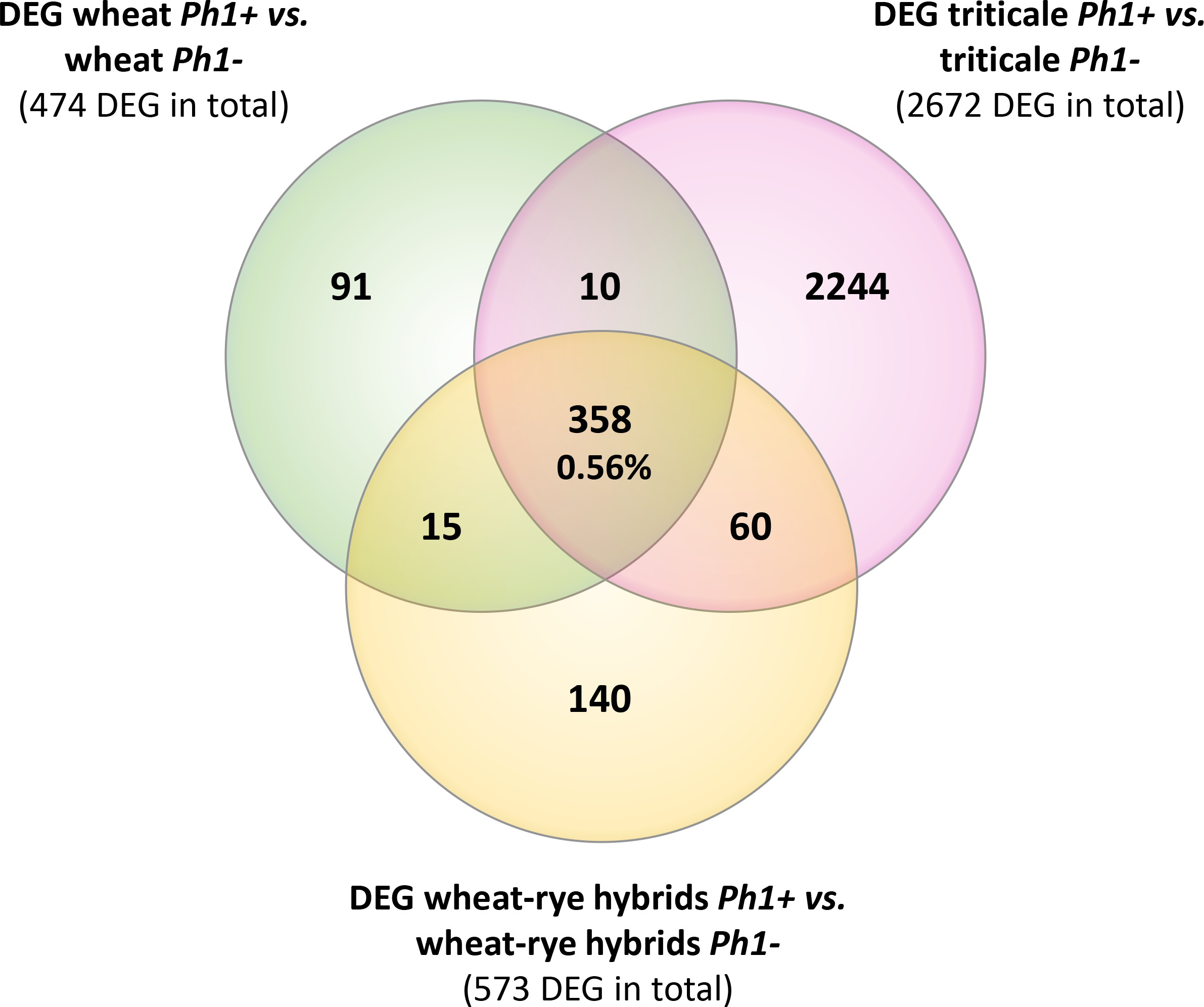
Venn diagram showing the overlapping DEGs common in all samples containing the *Ph1* locus (*Ph1+*) **vs.** all samples lacking it (*Ph1−*). Only 358 genes were differentially expressed in all genotypes lacking *Ph1*.

### 4. Genes responsible for the *Ph1* locus phenotype on recombination

In 1977 (Sears, 1977), the *Ph1b* deletion used in this study (and most studies involving this locus) was obtained and estimated to be of 70Mb in size. Using our gene expression data and the RefSeqv1.0 assembly, the *Ph1b* deletion is now defined to a 59.3 Mb region containing 1187 genes, from which 299 genes are expressed in our RNAseq data. The locus was further defined to a smaller region (Griffith et al., 2006; Al-Kaff et al., 2008), now defined to 0.5Mb in size and containing 25 genes (7 HC + 18 LC genes), from which only two are expressed in our RNA-Seq data. One of these two genes is the duplicated *ZIP4* gene (*TaZIP4-B2*), which has been recently identified as the gene responsible for both promoting homologous CO and restricting homoeologous CO (Rey et al., 2017, 2018a). Another gene within the *Ph1b* deletion, termed by the authors as *C-Ph1*, was also recently proposed to contribute to the *Ph1* effect on recombination, specifically during metaphase I (Bhullar et al., 2014); however, this gene does not show any expression in our RNA-Seq data. The authors reported 3 copies of this gene, one on 5A (truncated), one on 5B (with a splice variant named 5B^alt^) and one on 5D, claiming that only the 5B copy was metaphase I-specific and therefore, responsible for the phenotype characteristic of *Ph1*. However, blasting these gene sequences against the RefSeqv1.0 assembly showed that 5B^alt^ was a fourth gene copy located on 5A chromosome just upstream of the original 5A copy (5A-1: TraesCS5A01G381600LC and 5A-2: TraesCS5A01G381700LC). The RNA-Seq data obtained in the present study, as well as 849 wheat RNA-Seq samples now publicly available (Ramírez-González et al., 2018) and 8 wheat meiotic libraries available at https://urgi.versailles.inra.fr/files/RNASeqWheat/Meiosis/, can now be used to study the expression profile of all the different copies of the *ZIP4* and *C-Ph1* genes found across a diverse range of tissues and developmental stages (**Figure 4**). The TaZIP4 copy on 5B (*TaZIP4-B2*) is the dominant *ZIP4* copy and it is expressed in all tissue types, including all meiotic stages. *C-Ph1* has an almost tissue-specific expression pattern, limited to stamen tissue during the heading stage (post-meiosis stage), and with the 5D copy being expressed dominantly over all other gene copies (expression level of the 5D copy being >1700 TPM, and of the 5B copy being <31 TPM). None of the *C-Ph1* copies is expressed during any meiosis stage, except for the 5D copy that is expressed at a very low level (<4 TPM). In summary, we can confirm that *C-Ph1* on 5B is not expressed during meiosis, and cannot therefore be responsible for any *Ph1* effect during metaphase I.

**FIGURE 4.**
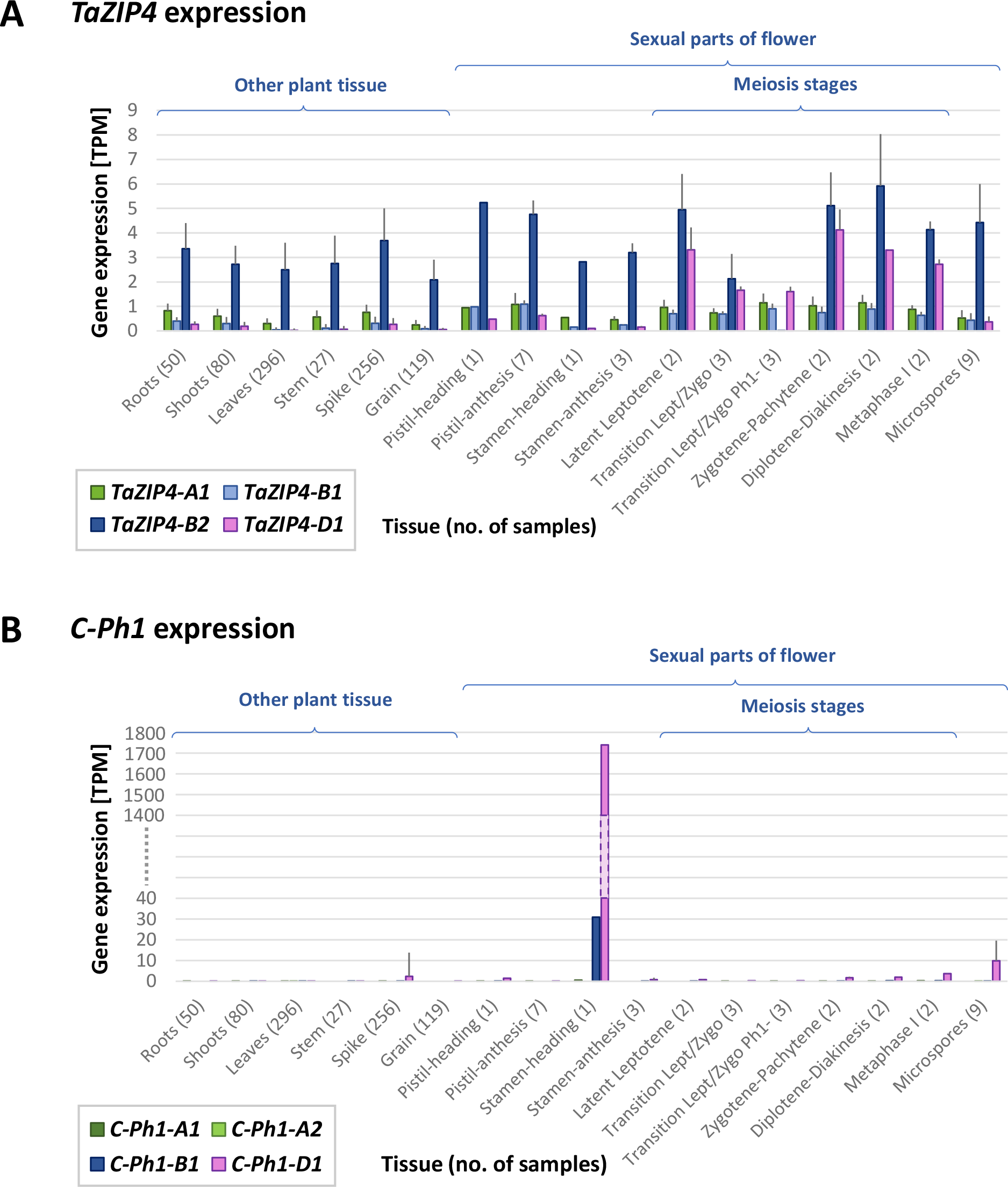
TaZIP4 and *C-Ph1* gene expression patterns across a range of tissues and developmental stages including meiosis. **(A)** *TaZIP4* copy on 5B (*TaZIP4-B2*) is the dominant *ZIP4* copy and it is expressed in all types of tissues, including all meiotic stages. **(B)** *C-Ph1* copy on 5B is not expressed during meiosis. The dominant *C-Ph1* copy is on 5D, which is mostly expressed during heading stage. 863 RNA-Seq samples were used to produce these graphs: 849 wheat RNA-Seq samples publicly available (Ramírez-González et al., 2018), eight meiotic samples available at https://urgi.versailles.inra.fr/files/RNASeqWheat/Meiosis/ and six samples from the present study. All RNA-Seq samples were mapped to the Chinese Spring RefSeqv1.0+UTR transcriptome reference and TPM values were calculated according to Ramírez-González et al., (2018). Error bar represents Standard Deviation value.

### 5. Cytological characterization of the newly synthesized triticale in the presence and absence of *Ph1*

Cytological analysis of wheat and wheat-rye hybrids, both in the presence and absence of the *Ph1* locus, has been well documented in previous studies (Orellana, 1985; Naranjo et al., 1988; Wang and Holm, 1988; Benavente et al., 1998; Mikhailova et al., 1998; Sánchez-Morán et al., 1999, 2001); however, this is the first time to our knowledge, that triticale lines lacking *Ph1* have been generated. As the chromosome coverage plots suggest the presence of several chromosome rearrangements, we performed mitotic and meiotic analysis to explore the origin and extent of these reorganizations.

#### 5.1 Chromosome configuration on mitotic metaphase cells

Root tip mitotic metaphase spreads were analysed by GISH (Genomic *in situ* hybridisation) to determine the extent of homoeologous recombination or any other rearrangements in the newly formed triticale lines, both containing and lacking *Ph1*.

Most of the triticale plants containing *Ph1* were euploid (six of nine plants), with a chromosome number of 56, and possessing 14 chromosomes from the A, B and D genome, plus 14 chromosomes from rye (**Table S8**). However, three of the triticale plants were aneuploid (**Figure 5A**), all with rye chromosomes missing. Interestingly, although some aneuploidy was observed, none of the triticale plants presented any inter-genomic translocations, suggesting that all recombination took place between homologous chromosomes. The only translocation observed in these triticales was the ancient translocation present in bread wheat T4A∙7B (**Figure 5A**).

**FIGURE 5.**
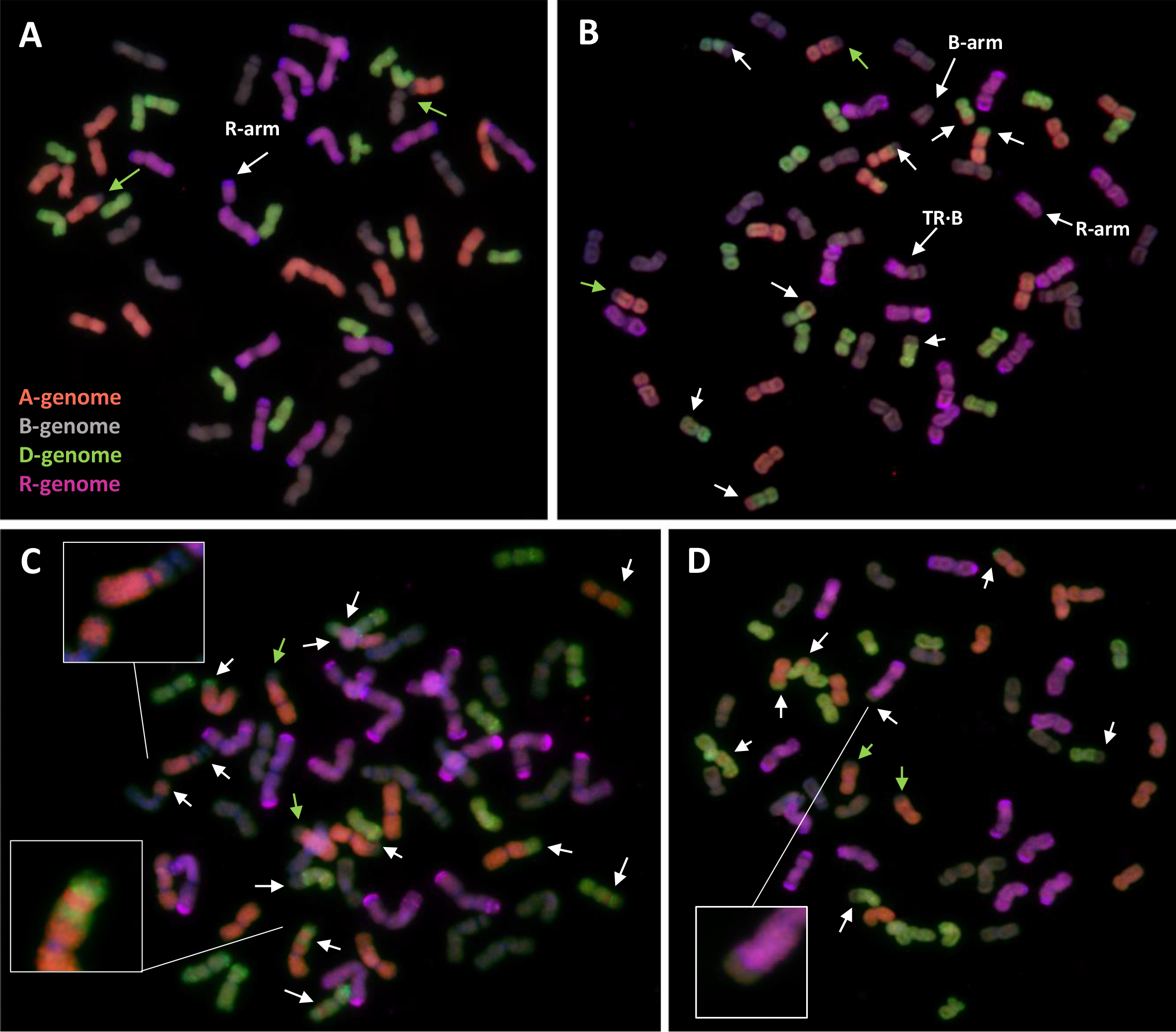
Root-tip metaphases of triticale analysed by GISH. **(A)** Triticale containing *Ph1* with 14 A-chromosomes, 14 B-chromosomes, 14 D-chromosomes and 13+arm rye (R) chromosomes (cn=55+arm). **(B)** Triticale lacking *Ph1* with 12 A, 14+arm B, 15 D, 10+arm R and a Robertsonian translocation between a rye and a B-chromosome (TR∙B) (cn=53 + 2arms). (C) Triticale lacking *Ph1* with 13 A, 15 B, 12 D and 14 R (cn=54). A proximal recombination between an A- and a B-chromosome, and an A-chromosome showing the result of three recombination events are highlighted. **(D)** Triticale lacking *Ph1* with 12 A, 13 B, 15 D and 13 R (cn=53). The result of recombination between a rye and a B-genome chromosome is highlighted. Reorganizations are indicated by white arrows. The ancient translocation T4A∙7B is indicated by green arrows.

In contrast to triticale containing *Ph1*, there were numerous chromosome rearrangements in the progeny of triticale lacking *Ph1*, including aneuploidy, deletions and intergenomic translocations. All individuals analyzed were aneuploids, with chromosome numbers ranging from 51 to 59 plus one chromosome arm (**Figure 5B-D**; **Table S8**). These triticale lines had only undergone three rounds of meiosis after synthesis, however some lines exhibited reorganizations corresponding to 16 recombination events between homoeologous chromosomes (**Table S8**). This wide range of chromosome rearrangements corresponds to the high levels of variability in the individual replicates of RNA-seq data shown in coverage plots (**Figure S4**). We even detected recombination between rye and a B-genome chromosome (**Figure 5D**), which is normally a very rare event. Apart from non-homologous recombination and aneuploidy, there were also other structural rearrangements, such as several individual chromosome arms and a Robertsonian translocation between a B-genome and a rye chromosome (**Figure 5C**). Recombination in cereals is normally restricted to the distal ends of the chromosomes, with 90% of wheat COs occurring in only 40% of the physical chromosome (Saintenac et al., 2009); interestingly, we also observed very proximal recombination events (**Figure 5C**), which is another example of the high level of reorganization present in these lines.

#### 5.2 Meiotic metaphase I configuration

Octoploid triticale have been reported to show meiotic instability and frequent aneuploidy in the presence of *Ph1*, resulting in reduced fertility (Scoles and Kaltsikes, 1974; Müntzing, 1979; Gustafson, 1982; Fominaya and Orellana, 1988; Lukaszewsky and Gustafson, 2011).

One third of plants in this study showed aneuploidy, in all cases involving rye chromosomes. Fertility rate was also reduced, even in euploid plants. All analyzed triticale lines containing *Ph1* showed a fairly normal meiosis with mostly bivalents being formed at metaphase I (**Figure 6A**); however, univalents were also frequently present (**Figure 6B**), as well as a low level of multivalents. GISH was performed on meiotic metaphase I cells to determine the origin of the univalents and to ascertain whether the bivalents were always between homologues. Most of the univalents observed were rye in origin, although some wheat origin univalent were also observed (**Figure 6C**). As for the bivalent formation, all were between chromosomes from the same genome (**Figure 6C**), suggesting that although there was some level of CO failure, no recombination between homeologues was taking place. This meiotic analysis supported data observed in the mitotic analysis and was consistent with the presence of *Ph1*. Although all the lines were fertile, the seed set was not complete in all flowers, and varied among different lines. This suggests that the abnormalities observed sometimes at meiosis, produced problems in chromosome segregation and probably aneuploidy and pollen abortion.

**FIGURE 6.**
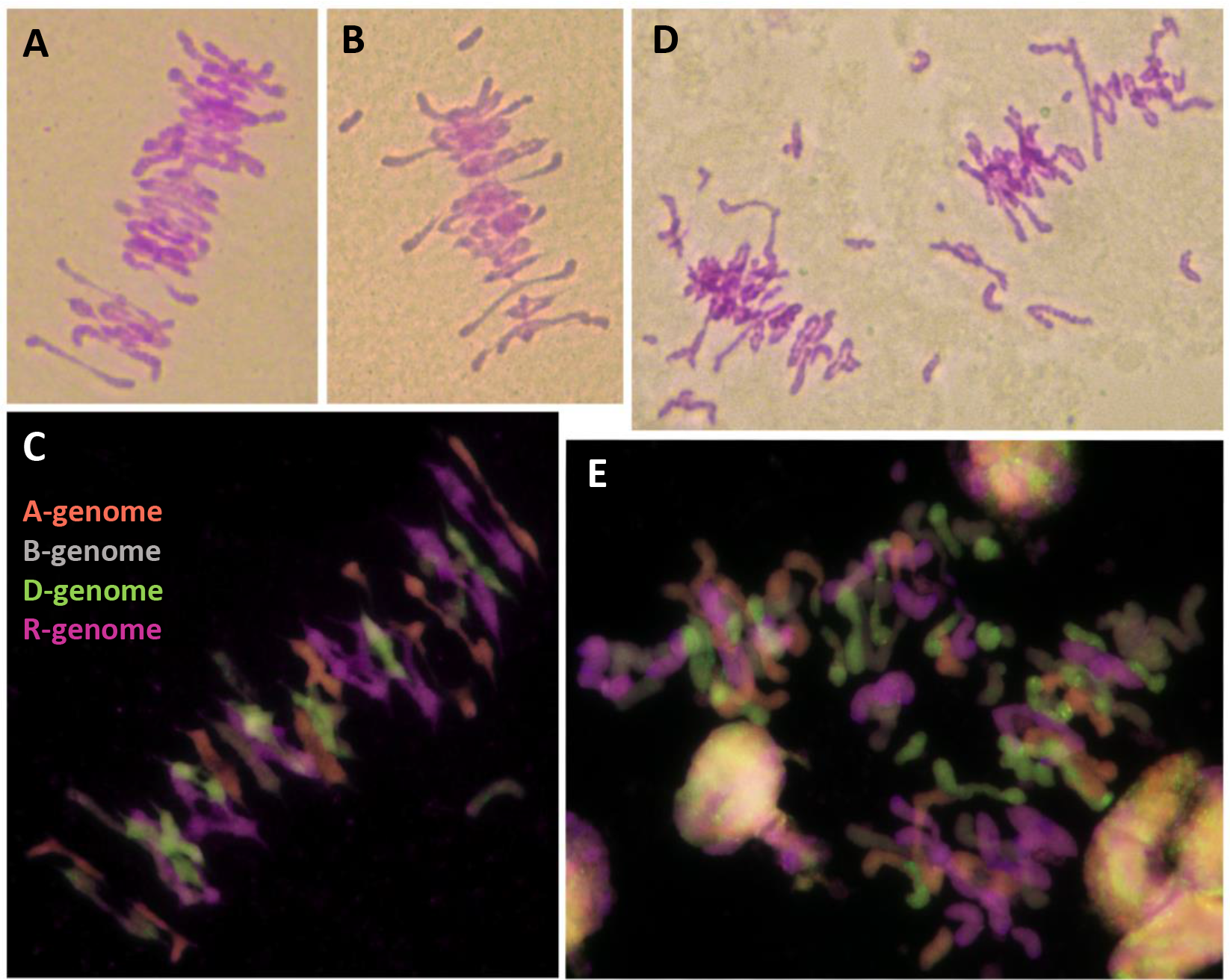
Meiotic metaphase I of triticale containing the *Ph1* locus (*Ph1+*) and lacking it (*Ph1*-), stained by Feulgen **(A, B, D)** and analysed by GISH **(C, E)**. **(A, B, C)** Triticale *Ph1*+ showing a fairly normal metaphase I. (D, E) Triticale *Ph1* - showing univalent, multivalents and chromosome fragmentation.

In the case of octoploid triticale lacking the *Ph1* locus, meiosis could not be analyzed in some plants because anthers had not developed properly. When meiotic metaphase I cells could be analyzed, substantial irregularities were observed, including univalents, multivalents and chromosome fragmentation (**Figure 6D**). GISH analysis showed that although most CO formation was between chromosomes from the same genome, chromosomes from all genomes were also involved in non-homologous association, particularly those derived from A- and D-genomes. Most fragmentation was of rye chromosomes (**Figure 6E**). These specific plants were all sterile apart from one plant, which produced 3 seeds. The fertility of all triticale lacking the *Ph1* locus used in this work was very low and decreased exponentially with each generation.

Morphology of all triticale plants containing *Ph1* was perfectly normal (**Figure S5**). However, every triticale plant lacking *Ph1* was morphologically different, with some exhibiting very abnormal phenotypes, likely to be the result of extensive chromosomal rearrangements (**Figure S5**).

## DISCUSSION

A high-quality annotated reference genome sequence of bread wheat (RefSeqv1.0) has recently been released by the International Wheat Genome Sequence Consortium (IWGSC, 2018), giving access to 110,790 high-confidence (HC) and 158,793 low-confidence (LC) genes. Together with this release, an extensive gene expression dataset of hexaploid wheat has been analyzed to produce a comprehensive, genome-wide analysis of homoeologue expression patterns in hexaploid wheat (Ramírez-González et al., 2018). In total, 850 available RNA-Seq data have been used across a diverse range of tissues, developmental stages, cultivars and environmental conditions. However, no specific data were available from meiosis, a key process ensuring proper chromosome segregation (and thus, genome stability and fertility) and leading to novel combinations of parental alleles, forming the basis of evolution and adaptation. This lack of meiotic RNA-Seq data is a general issue in many species, not only in wheat, due to the challenge of collecting plant material at specific meiotic stages. Some RNA-Seq approaches have been performed mainly in Arabidopsis, rice, maize, sunflower and brassica (Chen et al., 2010; Dukowic-Schulze and Chen, 2014; Dukowic-Schulze et al., 2014; Flórez-Zapata et al., 2014; Zhang et al., 2015; Braynen et al., 2017), but to our knowledge, no RNA-Seq analysis has previously been reported on wheat meiosis.

In this study, we took advantage of the recently released RefSeqv1.0 wheat assembly and our experience working on wheat meiosis, to perform an RNA-Seq analysis from six different genotypes: wheat, wheat-rye hybrids and newly synthesized triticale, both in the presence and absence of *Ph1*. All plant material was collected during early prophase, at the leptotene-zygotene transition, coinciding with telomere bouquet formation and synapsis between homologues. We addressed three questions in the study: whether overall wheat transcription was affected by the level of synapsis (and chromatin structure changes at the time of homologue recognition); whether wheat transcription was reshaped upon genome duplication; and whether wheat transcription was altered in the absence of the *Ph1* locus. Surprisingly, the answer to all three questions was negative. Wheat transcription was not affected in any of the three situations, revealing an unexpected level of transcription stability at this very important developmental stage, and suggesting a very important role (probably more than anticipated) for post-transcriptional regulation during meiotic prophase I. These results contrast with observations in somatic tissue of resynthesized hexaploid wheat, where 16% of genes were estimated to display nonadditive expression (Pumphrey et al., 2009).

### 1. High stability of global gene expression during early meiotic prophase

During meiosis, homologous (and sometimes non-homologous) chromosomes pair and then synapse through the polymerization of a protein structure known as the synaptonemal complex, which provides the structural framework for recombination to take place. Throughout the whole of meiosis, but particularly during the synaptic process, there are multiple changes in chromatin structure and organisation (and even positioning in the nucleus), taking place within a relatively short period of time and needing to be highly regulated. In wheat, synapsis is initiated during the telomere bouquet stage at early prophase, during the transition leptotene-zygotene (Martín et al., 2017). Moreover, it has been observed, that the process of recognition between homologues is associated with major changes in the chromatin structure of chromosomes (Prieto et al., 2004), suggesting that these changes in chromatin structure may be required for the homologue recognition process and initiation of synapsis. Indeed, it is now well understood that chromatin conformation is a critical factor in enabling many regulatory elements to perform their biological activity, and that chromatin structure profoundly influences gene expression (Dixon et al., 2015; Dogan et al., 2018). Therefore, it was reasonable to suppose that differences in synapsis, and therefore, in chromatin structure would translate into differences in transcription.

Surprisingly however, we did not find the expected differences in overall wheat transcription and gene expression when comparing samples with different levels of synapsis, indicating that the structural changes associated with this process were not directly coupled to transcription. Probably the clearest example of this was the comparison of wheat-rye hybrids and octoploid triticale. Wheat-rye hybrids possess a haploid set of wheat and rye chromosomes (there are no homologues present) and no synapsis is observed during the telomere bouquet stage (Martín et al., 2017). In contrast, octoploid triticale, which is obtained after chromosome doubling of wheat-rye hybrids, and which therefore possesses a whole set of wheat and rye chromosomes, exhibits extensive synapsis during the same stage. However, only 0.47% of the expressed genes were differentially expressed between these two samples (0.38% considering only HC genes), with most of these genes being involved in stress response and other metabolic processes (general functions) not related to meiosis.

Even more striking is the fact that in our study, global gene expression was not affected by whole genome duplication. Several previous studies have reported genetic and epigenomic processes being disrupted after hybridisation and polyploidisation, with subsequent changes in gene expression (Qi et al., 2012; Renny-Byfield et al., 2014; Khalil et al., 2015; Edger et al., 2017 and references therein). These studies were not performed on meiotic tissue, however given the known failure of meiosis in wheat-rye hybrids and the relatively normal meiotic progression in the duplicated triticale, it would have been reasonable to expect an effect on the expression pattern. However, as mentioned above, only a small fraction of genes were differentially expressed, again with none involved in meiosis. Unfortunately, the rye genome sequence is incomplete, and a similar analysis for rye genes could not be performed. In the future, when the complete rye genome sequence is available, it will be interesting to assess whether global expression of rye genes is also unchanged.

We propose two possible explanations for the striking robustness in gene expression during early meiotic prophase. The first possibility is that the transcription of genes required for the meiotic program has already occurred prior to the leptotene-zygotene transition. Meiosis is a very complex process which takes place in a relatively short period of time. In wheat, the whole meiosis process lasts only 24h at 20 °C, with the whole process of synapsis being less than 6h long (Bennet, 1973). Therefore, it would be reasonable to suppose that most of the transcription needed for such a critical process has already occurred prior to synapsis initiation. From mouse studies, it has been recently reported that a considerable number of genes involved in early, as well as later meiotic processes, are already active at early meiotic prophase (da Cruz et al., 2016). Moreover, a major change in gene expression patterns occurs during the middle of meiotic prophase (pachytene), when most genes related to spermiogenesis and sperm function appear already active (da Cruz et al., 2016). It is possible that this change in gene expression pattern also happens in wheat, with a transcriptional switch from pre-meiosis to meiosis taking place very early, before meiotic prophase. Only when more RNA-Seq datasets are available, can the dynamics of gene expression during wheat meiosis be fully understood.

A second possible explanation is that meiosis is mostly regulated at the post-transcriptional and post-translational level. There are several examples of post-transcriptional regulation during meiosis (Yelina et al., 2015 and references therein; Termolino et al., 2016 and references therein). Gene expression can be controlled at the RNA level, both quantitatively and qualitatively, and at various steps during RNA processing (alternative splicing, RNA editing, RNA silencing and translation regulation among others), with important consequences for the availability of different kinds of transcripts, and ultimately of proteins. The importance of the RNA interference (RNAi) machinery during meiosis has become clear in recent years (Dukowic-Schulze et al., 2014, 2016; Flórez-Zapata et al., 2016; Wu et al., 2017; Oliver et al., 2017), with small RNAs adding additional layers to the regulation of gene expression. A good example of this are phasiRNAs, a type of 21− and 24-nt phased, secondary siRNA from non-repeat regions produced specifically in male monocot meiotic transcriptomes (Johnson et al., 2009; Song et al., 2012), and whose function remains elusive. After translation, proteins can also acquire a great diversity of modifications, affecting their activity, localisation, turnover and interactions, as well as modifying and controlling their function. A plethora of proteins are interacting in a complex protein network during early prophase, and therefore, rapid modifications such as phosphorylation and dephosphorylation supply reversible means to regulate protein action (Josefsberg et al., 2002). Numerous post-translational modifications (PTMs) are known to take place during meiosis. Those involved in histone modifications (Sasaki et al., 2008; Xu et al., 2009; Kota et al., 2010; Greer et al., 2011; Yelina et al., 2014; Székvölgyi et al., 2015; Termolino et al., 2016) are frequently associated with changes in transcription, which is one of the reasons why we expected to detect larger changes in the meiotic transcription profile. However, several PTM are also involved in the modification of other meiotic proteins (Watanabe et al., 1997; Carballo et al., 2008; Fukuda et al., 2012; Sato-Carlton et al., 2017). Most of these meiotic PTMs have been reported in yeast and mammals, where synaptonemal complex formation and recombination are regulated through PTMs such as ubiquitination and phosphorylation. There are also similar reports of such modifications in plants (Lambing et al., 2018 and references therein). The results in the present study suggest that the role of post-transcriptional and post-translational modifications during wheat prophase is probably more important than initially thought, and that these modifications might be responsible for the control of this critical stage, rather than simple changes in the gene expression profile. In the future, more of these studies will be required in polyploid plants such as wheat, since the complexity of these genomes and their need to adjust their meiotic program to the presence of multiple genomes will only be understood by specifically studying these species directly.

### 2. Changes in wheat expression lacking the *Ph1* locus are the result of multiple chromosome reorganizations

The *Ph1* locus in wheat is by far, the best characterized locus involved in the diploid-like behaviour of polyploids during meiosis. Recently, the duplicated ZIP4 gene inside the *Ph1* locus on 5B (*TaZIP4-B2*) has been identified as responsible for the effect of this locus on homoeologous recombination (Rey et al., 2017, 2018a). ZIP4 is a meiotic gene shown to have a major effect on homologous COs in both *Arabidopsis* and rice (Chelysheva et al., 2007; Shen et al., 2012). Although its exact mode of action is unknown, it seems to act as a hub, facilitating physical interactions between components of the chromosome axis and the CO machinery (Perry et al., 2005; Tsubouchi et al., 2006). In diploid species, knockouts of this gene result in sterility, as failure of homologous COs at metaphase I leads to incorrect segregation. However, hexaploid wheat lacking *TaZIP4-B2*, only exhibits a small reduction in CO number, and still has fairly regular segregation. This is due to the four copies of *ZIP4* present in hexaploid wheat: one copy on each of the homoeologous group 3 chromosomes (*TaZIP4-A1, TaZIP4-B1 and TaZIP4-D1*) and a fourth copy on chromosome 5B (*TaZIP4-B2*). *TaZIP4-B2* is a transduplication of a chromosome 3B locus (IWGSC, 2018) and most probably appeared within the *Ph1* locus upon polyploidisation. In wheat lacking *Ph1* (and therefore *TaZIP4-B2*), ZIP4 copies on the homoeologous group 3 are still present, allowing CO formation, even if a small fraction occurs between non-homologous chromosomes. In the case of an allopolyploid species such as wheat, the process of homologue recognition is further complicated compared to diploids, by the presence of homoeologous chromosomes. We can hypothesise that upon polyploidisation, the newly formed hexaploid wheat already had mechanisms in place for the meiotic sorting of homologues from homoeologues during the telomere bouquet. This would have provided the new allopolyploid with some fertility until improved by the transduplication of *ZIP4* from 3B into the *Ph1* locus, with further modification and stabilisation of the meiotic process. Over time and evolution, the efficiency and stability of meiosis could be completely established. The transduplication of *ZIP4* is an example of how the meiotic program could be modified in polyploids in general, and wheat in particular, during evolution. It also illustrates the requirement to study such processes directly in these crop species, where there is a potential for manipulation in breeding programs.

*C-Ph1*, which is a syntenic ortholog of the RA8 gene in rice, has also been proposed to contribute to the *Ph1* effect on recombination (Bhullar et al., 2014). The RA8 gene in rice encodes an anther-specific BURP-domain protein expressed specifically in the tapetum, endothecium and connective tissue, but not in pollen grains, starting from the tetrad stage and reaching the maximum level of expression at the late vacuolated-pollen stage (Ding et al., 2009). Thus, *RA8* is suggested to play an important role in microspore development and dehiscence of anther (Jeon et al., 1999). Moreover, knockouts of this gene have been reported to induce male sterility (Patents WO2000026389 A3 and US20040060084). Expression profile analysis of all different copies of the *C-Ph1* gene using data generated in the present study, 849 wheat RNA-Seq samples now publicly available (Ramírez-González et al., 2018), and the meiotic RNA-Seq libraries deposited at https://urgi.versailles.inra.fr/files/RNASeqWheat/Meiosis/ reveals that the *C-Ph1* copy on 5D (rather than on 5B) is by far the most dominantly expressed. In addition, the newly generated wheat genome assembly reveals that the VIGS hairpin construct used for *C-Ph1* silencing (Bhullar et al., 2014), and which yielded sterility phenotypes, was designed from the wheat expressed sequence tag (EST) homolog BE498862 (448 bp), which is 100% identical to the 5D gene copy and not the 5B copy. As for the expression pattern, none of the *C-Ph1* copies show any meiotic expression, being mostly expressed afterwards during pollen formation, in common with the *C-Ph1* ortholog in rice RA8. These observations explain why our deletion covering the 5B copy of *C-Ph1* did not exhibit meiotic phenotype or sterility (Roberts et al., 1999; Al-Kaff et al. 2008), and suggest that *C-Ph1* is actually involved in microspore development and dehiscence of anther. Finally, *Ph1* is the dominant gene suppressing homoeologous CO within the wheat genome. Yet neither the presence of wild type *C-Ph1*, nor that of any other gene could suppress homoeologous CO induced by mutating *ZIP4* on 5B (*TaZIP4-B2*), being the same level of homoeologous CO to that observed in *Ph1b* deletion mutants (Rey et al., 2017, 2018a). We suggest that the use of the term *C-Ph1* for this gene is therefore misleading and should be replaced with a more appropriate description.

#### 2.1 Global gene expression during early prophase is not affected by Ph1

In the present study we used a total number of 21 different *Ph1b* mutant plants, to assess whether global gene expression was affected by the absence of this locus during early prophase. We identified a set of genes which were differentially expressed in all samples lacking *Ph1* compared to all samples containing *Ph1*. Only 358 genes were differentially expressed (0.56% of all expressed genes), of which 186 were located within the region corresponding to the 5B deletion, and 139 were located within the regions corresponding to the 3A and 3B deletions present in all *Ph1* samples. Therefore, no major global changes in wheat expression were observed in wheat lacking *Ph1* during early prophase. The effect of the *CDK2-like* genes inside the *Ph1* locus on premeiotic replication and the associated effects on chromatin and histone H1 phosphorylation did not subsequently affect overall gene expression in early meiotic prophase. Moreover, the significant structural changes observed in the absence of *Ph1* (centromere pairing and telomere dynamics during premeiosis, subtelomeric decondensation upon homologous recognition) were not associated with changes in global gene expression. This result is consistent with our previous conclusion that gene expression during meiotic early prophase is very stable and quite resilient to changes in chromatin structure.

#### 2.2 Wheat lacking Ph1 accumulate extensive chromosome rearrangements

The mean number of COs in wheat lacking *Ph1* (or *TaZIP4-B2*) was only 4-5 COs fewer than in wild type wheat (Martín et al., 2014; Rey et al., 2017). However, in the absence of *Ph1*, some COs can be formed between non-homologues, leading to non-homologous recombination and the accumulation of chromosome rearrangements, resulting in sterility after a certain number of generations. This karyotypic instability in the absence of *Ph1* has been previously reported (Sánchez-Morán et al., 2001), but the results obtained in the present work reveal that the intergenomic exchanges and deletions are higher than anticipated. All previous reports were based on GISH (Genomic in situ hybridisation) analysis, which only enables large rearrangements to be detected, mostly translocations, but no deletions. The present study, exploiting RNA-Seq analysis, has revealed both multiple translocations and deletions. The original *Ph1b* mutant was obtained in 1977 (Sears, 1977) and since then, intergenomic exchanges and other reorganizations have probably been accumulating. The present study reveals that as well as the *Ph1b* deletion on 5B, there are three further deletions in all *Ph1* mutant genotypes analysed. This suggests that they probably arose soon after the original *Ph1b* line was generated. Although our *Ph1b* mutant lines have been routinely backcrossed to wild type wheat after eight generations, extensive rearrangements still accumulate subsequently, meaning that every single *Ph1b* mutant could potentially be different. Thus, some of the effects previously attributed to the lack of the *Ph1* locus are likely to be the result of these reorganizations. However, if a sufficient number of different *Ph1b* mutant plants are used in any study, then this risk greatly decreases. In the present study, RNA seq samples were derived, and the data combined from 21 different *Ph1b* mutant plants. In the future, particularly for breeding purposes, we recommend the use of the *Tazip4-B2* TILLING mutant lines available at the UK Germplasm Resource Unit (https://seedstor.ac.uk/, code W10348 and W10349). These lines do not currently exhibit rearrangements, but will probably also accumulate reorganisations in further generations, so we recommend checking and cleaning the lines periodically.

#### 2.3 Extreme instability of triticale lacking Ph1

Octoploid triticale is the synthetic amphiploid resulting from the chromosome doubling of the hybrid between hexaploid wheat and rye. It is therefore, a new allopolyploid species. Primary octoploid triticale (containing *Ph1*) is unstable meiotically, with variable frequency of univalents in metaphase I and reduced fertility (Scoles and Kaltsikes, 1974; Müntzing 1979; Gustafson, 1982; Fominaya and Orellana, 1988; Lukaszewsky and Gustafson, 2011). For our RNA-seq analysis, triticale plants with 56 chromosomes were selected, ensuring that they all had the complete set of wheat and rye chromosomes. GISH analysis was performed on the progeny of plants used in the RNAseq experiments. This analysis revealed that one third of the progeny were aneuploids, with rye chromosomes always being the cause of aneuploidy. Interestingly, no recombination or reorganization between homoeologues was observed in any of the plants analysed, nor were any COs detected between homoeologues at meiotic metaphase I. This indicates that the origin of meiotic instability in octoploid triticale is most probably not related to the homologous recognition process, and that the presence of the *Ph1* locus in the wheat genome plays the same role in this new species, ensuring only homologous recombination. It has previously been speculated that late DNA replication of rye heterochromatin interferes with chromosome synapsis when rye chromosomes are placed in a wheat genetic background (Thomas and Kaltsikes, 1974, 1976; Merker, 1976); however, there are also reports contradicting this hypothesis (Fominaya and Orellana, 1988). There is a clear decrease in CO number in triticale compared to wheat and rye, as revealed by the frequent occurrence of univalents, particularly of rye chromosomes. Replication initiation activates a checkpoint system that prevents DSB formation in unreplicated DNA. Therefore, it is possible that late DNA replication of the terminal rye heterochomatic knobs prevents some DSB formation and/or affects the DSB repair pathway of these late breaks, preventing COs. This may explain the frequent presence of rye univalents. In the future, when the rye genome sequence is completed, it would be interesting to compare the expression of specific meiotic genes involved in recombination between wheat, rye and triticale, checking whether meiotic expression of both wheat and rye is altered when both genomes are placed together in the same cytoplasm (as a new species).

We also assessed the consequence on meiosis of generating triticale in the absence of *Ph1*, the locus responsible for the diploid-like behaviour of hexaploid wheat. As in the case of triticale containing *Ph1*, only plants with 56 chromosomes were selected for the RNA-Seq analysis. The fertility of these triticale plants lacking *Ph1* was extremely low, ranging from 9 seeds to complete sterility. GISH analysis on both somatic and meiotic cells showed that unlike in triticale containing *Ph1*, there were extensive reorganisations resulting from non-homologous recombination in the triticale lacking *Ph1*. Even though these triticale plants had only gone through three meiotic events since their synthesis, there were extensive recombination events between homoeologues. Chromosome fragmentation was also detected, particularly involving rye. There is a difference in heterochromatin DNA replication in the absence of the *Ph1* locus in wheat-rye hybrids (Greer et al., 2011). If, as has been suggested, late DNA replication of rye heterochromatin is the cause of triticale instability in the presence of *Ph1*, the additional delayed replication produced by the absence of *Ph1* could affect excessively DSB formation, causing not only a decrease in CO formation but also lack of DSB repair, and therefore, chromosome fragmentation. In any case, triticale lacking *Ph1* exhibits even more chromosomal rearrangements than wheat lacking *Ph1*, leading to an extreme phenotype and sterility.

### 3. Concluding remarks

Understanding polyploidization is of great importance in the understanding of crop domestication, speciation and plant evolution. One of the biggest challenges faced by a new polyploid is how to manage the correct recognition, synapsis and recombination of its multiple related chromosomes during meiosis, to produce balanced gametes. In the last few years, there has been a better understanding of the meiotic process from studies of diploid plants and other model organisms (Mercier et al., 2016). Polyploid crops have also benefitted from these advances since many of the key genes and processes seem to be conserved between species. However, polyploids differ considerably from diploids in many respects. Hexaploid wheat, with its large genome size, its high percentage of repetitive DNA and its three related ancestral genomes, is likely to have modified the meiotic process in adapting to its polyploidy. Surprisingly, here we found no evidence for major changes in gene expression during early meiotic prophase, despite variations in synapsis, whole genome duplication or the absence of the *Ph1* locus. This suggests that the transcription of genes required for early meiotic prophase has already occurred prior to this stage, and/or that many of the meiotic prophase events are regulated post-translationally. Genetic studies in polyploids such as wheat have lagged far behind diploid species, partly because of the lack of key genetic resources. However, the release of the RefSeqv1.0 assembly in hexaploid wheat (IWGSC, 2018), the availability of expression data, (including that generated from the present study) presented in a browser www.wheat-expression.com (Borrill et al., 2016; Ramírez-González et al., 2018) enabling easy visualization and comparison of transcriptome data, and the availability of TILLING mutants for every wheat gene (Krasileva et al., 2017), will now allow more rapid progress to be made in our understanding of meiosis.

## Acknowledgments

This work was supported by the UK Biotechnology and Biological Research Council (BBSRC), through a grant part of the Designing Future Wheat (DFW) Institute Strategic Programme (BB/P016855/1) and Grant BB/J007188/1. Next-generation sequencing and library construction were delivered via the BBSRC National Capability in Genomics (BB/CCG1720/1) at Earlham Institute by members of the Genomics Pipelines Group.

## Author contributions

AM, PS, and GM: conceived and designed the study; AM: obtained the hybrids and triticale, produced the meiotic RNA samples, did the cytological analyses, interpreted the results and wrote the first draft of the manuscript; PB carried out the differential expression analysis and assisted with analysis of all data. JH did the mapping using HISAT and created the chromosome coverage plots. AA: carried out some data analysis and assisted with analysis of data. RR: carried out the mapping using kallisto for the differential expression analysis and integrated the wheat analysis into the expVIP platform; PS and GM: contributed to the interpretation of the data; GM: edited the manuscript. All authors have read and approved the final version of the manuscript.

## Supporting Information

**TABLE S1** Number of cleaned reads generated and mapped for each sample. The RNA-Seq data were aligned to the RefSeqv1.0 assembly using HISAT with strict mapping options to reduce the noise caused by reads mapping to the incorrect regions.

**TABLE S2** Number of reads generated and mapped for each sample using Kallisto. **(A)** Wheat samples were pseudoaligned against the Chinese Spring RefSeqv1.0+UTR transcriptome reference. **(B)** Wheat-rye hybrids and triticale samples were pseudoaligned against a wheat+rye transcriptome constructed in silico by combining the Chinese Spring RefSeqv1.0+UTR transcriptome reference with the published rye transcriptome (Bauer et al., 2016).

**TABLE S3** Number of DEG among samples using different thresholds in the presence (*Ph1*+) and absence (*Ph1*-) of the *Ph1* locus. **(A)** Number of wheat DEGs. **(B)** Number of rye DEGs. The first number in every column tittle represents the p-adj filter (⍰ 0.05, ⍰ 0.01 or ⍰ 0.001). The 2FC indicates that genes were up or down-regulated over 2-folds.

**TABLE S4** Genes differentially expressed among samples and total number of expressed genes (EG) in this study. **(A)** High confidence (HC) wheat genes. **(B)** Low confidence (LC) wheat genes. **(C)** Rye genes.

**TABLE S5 (A)** Gene ontology (GO) classification of DEGs up-regulated in wheat-rye hybrids vs. triticale (both containing the *Ph1* locus). **(B)** Gene ontology (GO) classification of DEGs down-regulated in wheat-rye hybrids *vs*. triticale (both containing the *Ph1* locus). (C) GO Slim classification of DEGs down-regulated in wheat-rye hybrids vs. triticale (both containing the *Ph1* locus). This list was created to have a broad overview of the ontology content without the detail of the specific fine-grained terms.

**TABLE S6 (A)** Functional annotation of DEGs up-regulated in wheat-rye hybrids vs. Triticale (both containing *Ph1*). **(B)** Functional annotation of DEGs down-regulated in wheat-rye hybrids vs. triticale (both containing *Ph1*).

**TABLE S7** Functional annotation of the 33 DEGs shared by all samples lacking *Ph1 vs*. all samples containing *Ph1*, and which are not located in any of the common deletions present in all samples lacking *Ph1*.

**TABLE S8** Chromosome configuration of nine newly synthesized triticale both containing the *Ph1* locus (*Ph1+*) and lacking it (*Ph1*-). No inter-genomic recombination was detected in the presence of *Ph1*.

**FIGURE S1**. Principal Component Analysis (PCA) of samples analysed in this study. Three biological replicates were produced per genotype. **(A)** PCA for the six wheat samples, three containing the *Ph1* locus (*Ph1*+) and three lacking it (*Ph1*-). The x and y axis represent the two principal components of the total variance, 73% and 12% respectively. **(B)** PCA for 12 wheat-rye hybrid and triticale samples. Three hybrids containing and three lacking *Ph1*, and three triticale containing and three lacking *Ph1*. The x and y axis represent the two principal components of the total variance, 70% and 15% respectively.

**FIGURE S2**. Representation of the ratio of coverage along all chromosomes using Box plots. **(A)** Box plot comparing wheat-rye hybrids and triticale, both containing the *Ph1* locus. **(B)** Box plot comparing wheat in the presence and absence of *Ph1*. **(C)** Box plot comparing wheat-rye in the presence and absence of *Ph1*. **(D)** Box plot comparing triticale in the presence and absence of *Ph1*. Arithmetic mean values of the coverage ratio per chromosome are indicated on the upper part of the plots. Mean values ⍰ 0.05 and ⍰ - 0.05 are highlighted in magenta.

**FIGURE S3** Chromosome coverage plots of wheat containing *Ph1* (*Ph1+) vs.* each individual sample of wheat lacking *Ph1* (*Ph1*-). Heatmaps show that each wheat sample lacking *Ph1* is different. Several deletions (visualized in dark blue) are common to all three samples, but other deletions and chromosomes rearrangements are different between them.

**FIGURE S4** Chromosome coverage plots of triticale containing *Ph1* (*Ph1+) vs*. each individual sample of triticale lacking *Ph1* (*Ph1*-). Heatmaps show that each triticale sample lacking *Ph1* is different. Several deletions (visualized in dark blue) are common to all three samples, but other deletions and chromosomes rearrangements are different between them.

**FIGURE S5** Morphology of whole plants (A) and spikes (B) of triticale containing the *Ph1* locus (*Ph1+*) and lacking it (*Ph1*-). Plant and spike morphology of all triticale containing *Ph1* was perfectly normal; however, every triticale lacking *Ph1*, was morphologically different, some exhibiting very abnormal phenotypes.

